# Prioritizing persistent microbiome members in the common bean rhizosphere: an integrated analysis of space, time, and plant genotype

**DOI:** 10.1101/727461

**Authors:** Nejc Stopnisek, Ashley Shade

## Abstract

The full potential of managing microbial communities to support plant health is yet-unrealized, in part because it remains difficult to ascertain which members are most important for the plant. However, microbes that consistently associate with a plant species across varied field conditions and over plant development likely engage with the host or host environment. Here, we applied abundance-occupancy concepts from macroecology to quantify the core membership of bacterial/archaeal and fungal communities in the rhizosphere of the common bean (*Phaseolus vulgaris*). Our study investigated the microbiome membership that persisted over multiple dimensions important for plant agriculture, including major growing regions, plant development, annual plantings, and divergent genotypes, and also included re-analysis of public data. We found 48 core bacterial taxa that were consistently detected in all samples, inclusive of all datasets and dimensions. This suggests reliable enrichment of these taxa to the plant environment and time-independence of their association with the plant. More generally, this work provides a robust approach for systematically prioritizing core microbiome memberships in any host or system.

## Introduction

Agriculture requires more efficient use of available resources, and the naturally occurring, soil-dwelling microbiota offers potential to contribute to the responsible intensification of agriculture. Selection and breeding of plants for their beneficial associations with microbiota has promise to deliver a new generation of microbe-improved plants [1–6]. The ideal outcome of such efforts would achieve a balance of sustainable agriculture with food security. To achieve this, we must understand the relationships between plants and their associated microbiomes, including the differentiation of key or “core” members that engage with the plant directly from transient or opportunistic members that do not.

Common bean (*Phaseolus vulgaris* L.) is the most important food legume grown worldwide, and especially for developing economies in South America, Africa and Asia [7]. The origin of common bean is central Mexico, and from there it spread to central and to south America around 165,000 years ago [8]. This resulted in the development of two major and eco-geographically distinct common bean gene pools with partial reproductive isolation [9–11]. The Mesoamerican gene pool was distributed from northern Mexico to Colombia and the Andean gene pool ranged from southern Peru to northwestern Argentina. Since 8,000 years ago, each pool was separately and selectively bred, leading to further diversification between them [8, 12, 13]. Because of pre-existing genetic differences in each gene pool followed by divergent breeding history, common bean presently offers a distinctive opportunity for understanding how the host and the environment contribute to rhizosphere microbiome assembly.

The objective of this study was to apply approaches from macroecology to prioritize the persistent members of a core bacterial, archaeal, and fungal root microbiome inclusive of multiple gradients and categories of drivers expected to be important for plant agriculture. With the cooperation of the U.S. Bean coordinated agricultural project (Bean CAP), we executed a first-of-its kind study of two divergent bean genotypes grown under field conditions in five major North American bean growing regions, including Michigan, Nebraska, Colorado and Washington. These two genotypes belong to the Mesoamerican (Eclipse genotype) and Andean (California Early Light Red Kidney, CELRK genotype) gene pools that represent the major divergences from the wild bean ancestor [8]. For these two divergent genotypes, we also assessed how core members of the root microbiome changed over plant development and root compartment. We also used public data to perform a comparative analysis of the bacterial microbiome found in our study with microbiome members detected in other bean genotypes grown in South America [14, 15]. From our effort that was inclusive of both broad biogeography (including the U.S. and Colombia), plant development, and inter-annual plantings, we discovered a core bean rhizosphere microbiome of 48 members that persistently associated with this nutritionally, agronomically, and economically important crop. This core was discovered in spite of apparent microbiome differences that were attributable to local soil conditions and management. However, we did not detect an influence of plant genotype, suggesting that this core membership supersedes it.

## Material and Methods

### Study design, sampling and soil physicochemical analysis

We designed a biogeography study of two divergent bean genotypes (from Mesoamerican and Andean gene pools [16, 17]), both grown in the field in the summer of 2017 at five research and extension farms that represent major U.S. bean production regions (**Table 1**) [18]. The research and extension farms were: Saginaw Valley (SVERC), Michigan (MI), Montcalm county (MRF), MI, Scott Bluff county, Nebraska (NE), Fort Collins, Colorado (CO), and Othello, Washington (WA).

**Table 1:** Description of geographical, managemental and soil properties of the common bean growing locations included in the study.

| Sample same | Growing location | Elevation | Bean genotype | Weather (T, precipitation)* | Rotation history | Fertilization (per acre) | Irrigation | pH | Nitrogen (%) | Organic matter (%) |
| --- | --- | --- | --- | --- | --- | --- | --- | --- | --- | --- |
| SVREC | Saginaw Valley Research and Extension Center, MI, US | 190 m | CELRK | 24.5°C, 5.7 mm | Common bean, wheat, maize | Synthetic (400lbs of 15N-5 P-13K) | None | 7.7 (± 0.1) | 0.13 (± 0.00) | 2.33 (± 0.06) |
| MRC | Montcalm Research Center, MI, US | 280 m | Eclipse | 24°C, 7.4 mm | Common bean, maize, potato | Synthetic (200lbs of 19N-0P-19K) | Yes | 5.9 (± 0.4) | 0.10 (± 0.01) | 2.13 (± 0.06) |
| NE | Scottsbluff, NE, US | 1200 m | CELRK and Eclipse | 27°C, 2.18 mm | Common bean, maize, common bean | None | Yes | 7.9 (± 0.1) | 0.07 (± 0.01) | 1.39 (± 0.18) |
| CO | ARDEC, Fort Collins, CO, US | 1536 m | CELRK and Eclipse | 31°C, 1.4 mm ** | Common bean, maize, barley | Organic (220lbs urea (2016), 3 tons manure (2015)) | Yes | 8.1 (± 0.2) | 0.11 (± 0.00) | 1.7 (± 0.15) |
| WA | WSU, Othello, WA, US | 320 m | CELRK and Eclipse | 26.3°C, 0.2 mm | Common bean, maize, wheat | Synthetic (40lbs N/20lbs P/10lbs ZN2O5) | Yes | 6.1 (± 0.4) | 0.08 (± 0.01) | 1.78 (± 0.10) |

\* Temperature (as max air temperature) and precipitation are averaged for the period between May 1<sup>st</sup> and July 31<sup>st</sup> 2017.
\*\* Data available only for the period between Jun 28<sup>th</sup> and Jul 31<sup>st</sup> 2017.

Triplicate bean plants were grown in each of three (MI, WA, CO), or four (NE) plots, totaling 9 or 12 plants per growing location. Plants were harvested at flowering stage (mid to late July) because we wanted to analyze mature microbial communities and control for potential differences in the microbiome over early plant development.

We selected two common bean cultivars with very distinct genotypes: Eclipse [16] of Mesoamerican origin and California Early Light Red Kidney (CELRK), an old kidney bean landrace of Andean origin [17]. Even though we included only two bean genotypes, we selected highly divergent genotypes from far ends of the spectrum of the common bean genetic diversity; these lineages diversified after a biogeographic bottleneck that separated them between Central and South America [19]. We hypothesized that if there was an effect of bean genotype on the root microbiome, it should be measurable when comparing these genotypes from divergent lineages. Both bean genotypes were grown in each growing location, except in Michigan, where CELRK was grown at Saginaw Valley and Eclipse at Montcalm county.

Plant roots with attached soil were packed into bags, stored on ice and shipped immediately to Michigan State University for processing. From each sample, root-attached soil was collected by gently shaking the roots and designated as rhizosphere soil; unattached root-zone soil was also collected and used as bulk soil. It was expected that the rhizosphere soil contains a subset of microbiome diversity that is recruited from the bulk soil [20] but we note that these soils were not “bulk” in the traditional definition, in that they were collected from the agricultural fields and expected to be proximal to and influenced by the roots of previous crops. Soils were sieved through 4 mm sieves to remove any plant debris and larger soil minerals before soil physical-chemical analysis. Soil analysis was done at the Michigan State Soil Plant and Nutrient Laboratory on root-zone soil samples pooled by plant genotype and plot (see Supporting Text for details).

We designed a second temporal study to assess the dynamics of the core taxa over plant development. The two bean genotypes, CELRK and Eclipse, were grown at the same two sites in Michigan, U.S., Montcalm county and Saginaw Valley, in the summer of 2018. Plants root systems were harvested at 5 growth stages, including stage 1: V2 (appearance of second trifoliate), stage 2: V5 (appearance of fifth trifoliate), stage 3: flowering, stage 4: pod filling, and stage 5: senescence/drying). At each sampling time, roots were collected and transported on ice to the laboratory for immediate processing. Rhizosphere soil was collected by gently shaking roots, as described above. Any remaining, tightly-attached soil was designated as rhizoplane soil, and was collected by first vortexing roots in 1x phosphate buffer solution (PBS) for 4 min and finally removing the supernatant after centrifugation for 10 min at 8000 *g* and 4°C. Collected soil was immediately frozen in liquid nitrogen and stored at −80°C.

### Microbiome sequencing and analysis

There were 31 rhizosphere (attached to the root and pooled by plot) and 8 root-zone/bulk (one pooled sample per growing location and plant genotype) soil samples sequenced for microbiome analysis from the biogeography study, and 125 rhizosphere and 127 rhizoplane soil (individual plants) samples sequenced from the plant development study. DNA extractions, including negative controls, were performed as per standard soil protocols (see Supporting Text for details). 16S rRNA gene amplicon and ITS amplicon sequencing was performed at the Michigan State Genomics Core Research Support Facility. We processed the 16S rRNA gene amplicons using an open-reference clustering first against the SILVA v128 database [21, 22]. First, raw reads were merged, quality filtered, dereplicated, and clustered into 97% identity operational taxonomic units (OTUs) using the UPARSE pipeline (version 11, [23]). Reads not matching to the SILVA database were used for de-novo clustering at 97% sequence identity. Reference picked and de-novo reads were combined before taxonomy was assigned [21]. Taxonomic annotations for 16S rRNA gene OTU representative sequences were assigned in the QIIME 1.19 environment [24] using SILVA database [22]. ITS OTU representative sequences were taxonomically annotated using the CONSTAX tool [25] with the UNITE database version 7.2 [26]. OTUs with unassigned taxonomy at the domain level and OTUs annotated as mitochondria or chloroplasts were removed. Contaminant OTUs were removed using decontam package in R [27]. Additionally, we performed zero-radios OTU analysis (aka ZOTUs that are clustered at 100% sequence identity) for reads detected within the core OTUs. In short, first we subset all reads that were included within the core OTUs and processed them through the UNOISE3 pipeline [28] to create ZOTUs. Then we determined each ZOTU’s affiliation to their originating core OTU by matching the read identities between ZOTU and OTU clusters using a customized code (see GitHub repository).

Statistical analysis, including abundance-occupancy analysis to detect a core microbiome [29], and data visualization were performed in R (see Supporting Text). The co-occurrence network and global network properties were calculated using the Molecular Ecological Network Analysis Pipeline (MENAP) [30] and visualized with Cytoscape v.3.5.1 [31]. Comparative analyses with published datasets from Pérez-Jaramillo et al. [15] (BioProject ID PRJEB26084). was performed by downloading study raw reads from NCBI and processing as per our data pipeline to analyze both sets identically. An expanded description of all ecological statistics, MENAP parameters, and our combined core microbiome analysis with public data are provided in Supporting Text.

### Data and code availability

The raw sequence data are deposited in the NCBI Sequence Read Archive (BioProject ID PRJNA524532). All read processing steps, bioinformatic workflows, R code, and custom scripts are available on GitHub (https://github.com/ShadeLab/PAPER_Stopnisek_2019_BeanBiogeography).

## Results

### Sequencing summary and microbial diversity across growing regions

There were 31,255 to 506,166 and 22,716 to 252,810 reads per sample for 16S rRNA and ITS biogeography datasets, respectively. We rarefied samples to 31,255 reads for 16S rRNA gene amplicons and to 22,716 for ITS. With these thresholds, we achieved richness asymptotes for both datasets, suggesting that sequencing efforts were sufficient to capture comparative dynamics and diversity (**Fig. S1**). The total richness observed at this rarefaction depth was 1,505 fungal and 23,872 bacterial and archaeal OTUs.

As reported in other rhizosphere studies, the total fungal diversity was lower than bacterial /archaeal diversity in the rhizosphere of the common bean [32–34]. Richness varied by growing location (ANOVA, F value=12.4, *p*-value<0.0001 and F value=13.1, *p*-value<0.0001 for 16S rRNA and ITS data, respectively, **Fig. S2**) but was highest at the Montcalm Research Farm (Michigan, US) for both, bacteria/archaea and fungi. An analysis of community beta diversity revealed strong biogeographic patterns in community structure explained by location, soil pH and fertilization, in agreement with other literature [35–38] (**Fig. S3**, see **Supplemental Text**). However, an influence of plant genotype was either weak or not detected, which we partially attribute to plant growth in field conditions, following observations made in other plant species grown under field conditions [35, 36, 39] (**Fig. S2BD**, see **Supplemental Text).**

The common bean rhizosphere microbiome included the major expected lineages for both bacteria and fungi (**Fig. S4AB**), in agreement with other plant rhizosphere studies [40–45]. Together with previous studies, these data provide more evidence that root-associated microbial taxa are phylogenetically and potentially functionally conserved [46]. Proteobacteria, Acidobacteria, Bacteroidetes, and Actinobacteria collectively comprised on average 73.5% of the bacteria/archaeal community, Ascomycota dominated the fungal community with a mean total relative abundance of 53% with notable sample-to-sample variance (range from 16.5% to 84.5%).

### A core rhizosphere microbiome is detected across U.S. bean growing regions

We noticed a large number of OTUs that were shared among all growing locations them for the bacterial/archaeal communities (2,173 taxa, mean 31.5%, range 29.5% to 34.7%). There was a smaller but notable overlap for the fungal communities (70 taxa or mean 4.5%, range from 0.9% to 17.9%; **Fig. S4CD**). These data suggested that, despite measured edaphic differences across growing locations and strong biogeographic signal, the common bean rhizosphere recruited many similar taxa that could be functionally important for the bean. Therefore, we explored abundance-occupancy distributions of taxa ([47, 48] and references therein) to infer the core bean microbiome of taxa with an occupancy of 1 (i.e., found in all soil samples, all plots and across all growing locations; **Fig. 1AB**). Among bacteria and archaea, 258 phylogenetically diverse taxa were cosmopolitan in the dataset (**Fig. 1C**), including numerous and abundant Proteobacteria (117 OTUs) with a dominant taxon classified as *Arthrobacter* sp. (FM209319.1.1474, mean relative abundance of 1.43%). The bacterial/archaeal core also contained taxa of interest for potential plant benefits (e.g. *Sphingomonas, Rhizobium, Bacillus, Streptomyces*), as well as some genera that can be associated with disease (e.g. *Ralstonia*). There were 13 taxa in the fungal core (**Fig. 1D**), and these were largely composed of Ascomycota (10 OTUs), with dominating taxon OTU823 from the *Phaeosphaeriaceae* family (mean relative abundance 10.1%). Notably, taxa that were unique to either bean genotype were relatively rare and inconsistently detected (**Fig. 1**, orange and black points). Together, these results suggest that common bean consistently recruits particular microbiome taxa.

**Fig. 1:**
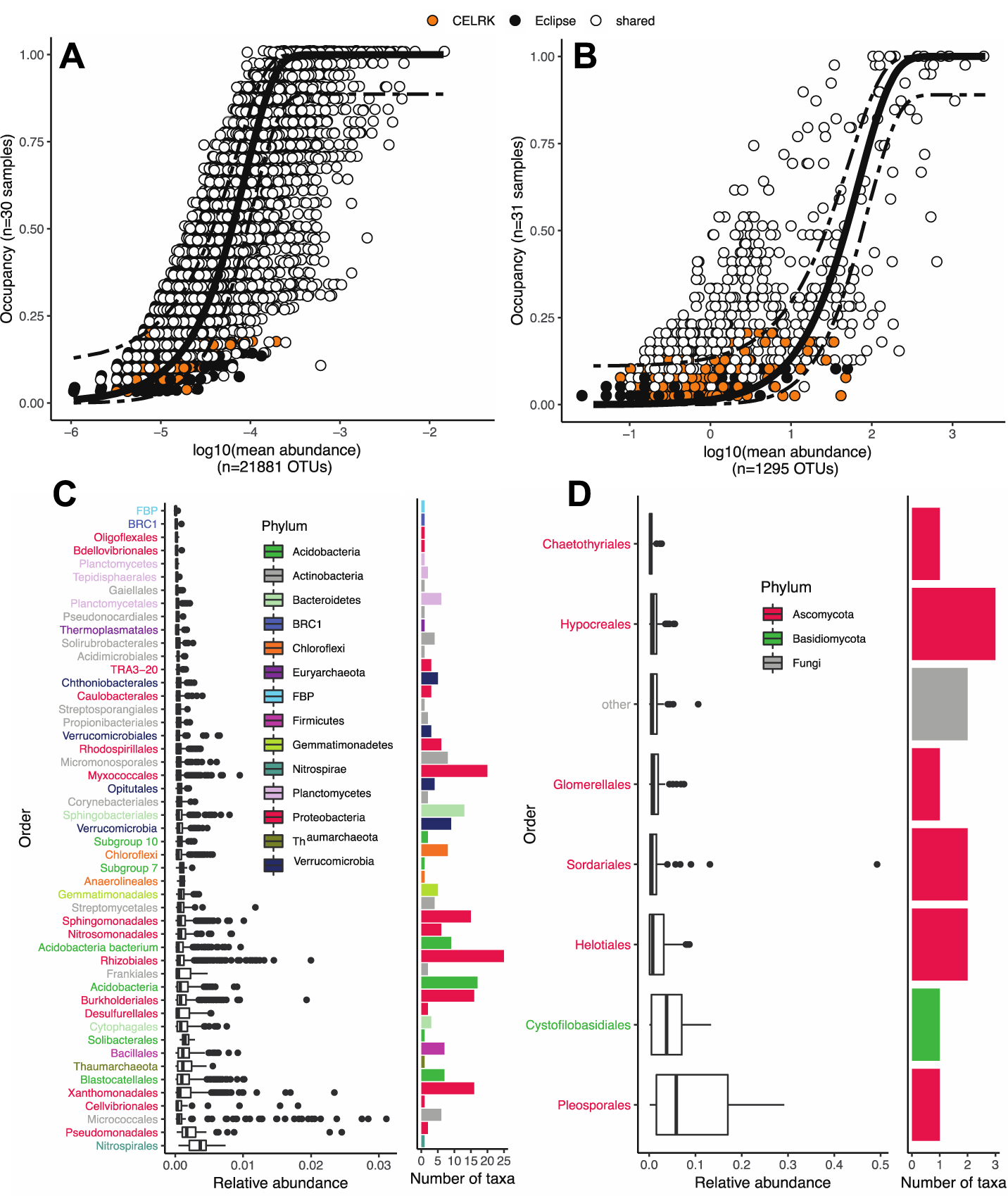
Abundance-occupancy distributions were used to identify core members of the rhizosphere microbiome for bacteria/archaea (A) and fungi (B). Taxa exclusive to a genotype are indicated in orange (CELRK) or black (Eclipse), and taxa shared across both genotypes are white. The solid line represents the fit of the neutral model, and the dashed line is 95% confidence around the model prediction. Taxa with an occupancy of 1 (i.e., detected in all samples) were considered members of the core. Relative abundance of these taxa is represented as boxplots, grouped by order and number of taxa therein (C, D). Panels C and D are color-coded by phyla level.

Next we wanted to investigate if the prioritized core taxa are indeed selected by the plant environment or assembled through neutral processes by applying the Sloan neutral model [49, 50]. The neutral expectation of abundance-occupancy distributions is that very abundant taxa will have high occupancy, while rare taxa will have low [48–51]. Taxa that deviate from the neutral expectation are more strongly influenced by deterministic factors, like environmental conditions, than by stochastic factors, like drift and dispersal. The neutral model fit of the abundance-occupancy distribution (solid line, **Fig. 1AB**) identified several taxa that had frequencies either above or below the 95% confidence intervals of the model (dashed lines). Specifically, 13.7% of the bacterial/archaeal and 30.4% of fungal taxa, deviated from the neutral expectation (**Table S3**). One hundred and seventy-one core taxa were predicted above the neutral model partition; these deterministically-selected taxa are prime candidates for follow-up studies of interactions with the host plant. Overall, the bacteria/archaea community had better fit to the neutral expectation than fungal (R^2^ of 0.74 and 0.34, and migration rates (m) of 0.301 and 0.003, respectively), suggesting that dispersal was relatively less limiting for the bacteria/archaea than for the fungi. This finding agrees with other work suggesting that fungi are more sensitive to local climate or more dispersal limited than bacteria [52–55].

### A core rhizosphere microbiome is detected for common bean grown on different continents

We wanted to better understand if these U.S. core taxa were associated with the bean rhizosphere across a larger geographical scale, which would suggest the potential for selective plant recruitment and cosmopolitan distribution of core taxa. Therefore, we compared our U.S. data to a recently published study of rhizosphere bacteria and archaea from common beans grown in Colombian agricultural soil [14]. The Colombian study offered a key contrast because it included eight divergent bean lineages, including wild (n=2), landrace (n=1), and cultivated genotypes (n=5), grown in soil from a different continent that has starkly different climate and management from the U.S. growing regions. To enable direct comparison, we re-analyzed raw reads and compared the datasets by matching to either the same taxon identifiers when clustered to SILVA database, or 100% identity by BLAST to *de novo* clustered reads (see Materials and Methods). Surprisingly, 39.6% (3,359 OTUs) of rhizosphere taxa from the Colombian-grown beans were also shared with the U.S. dataset (**Fig. 2**). Both datasets included taxa that were highly represented in the other: 62% of U.S. core (159 out of 258) were found also in Colombia, and 51% of Colombian core (433 out of 848) were shared with the U.S. (**Fig. 2A**). Core taxa were again defined stringently with an occupancy of 1, and 48 taxa were found across all samples, inclusive of both datasets. We refer to this as the “global” core to distinguish the subset from the larger group of core taxa inclusive to the US only (though note that this descriptor is for simplicity and that this does not include global representation of bean root samples on Earth). These global core taxa were composed of many Proteobacteria, with *Rhizobiales* showing the most consistent relative abundance between the studies (**Fig. 2B**, e.g. 0.187% and 0.138% in Colombia and U.S. dataset, respectively). Notably, none of this global core taxa were universally detected in very high abundance, and all but two OTUs (a U.S. *Arthrobacter* sp. and a Colombian *Austrofundulus limnaeus* with mean relative abundances of 1.43% and 1.01%, respectively) would be classified as rare by a typical mean relative abundance threshold below 1%, hinting to a potential role of rare taxa in providing key functions. A similar observation was made with rhizosphere microbiota of 19 herbaceous plant species, in which taxa of low abundance were among those significantly enriched in the rhizosphere as compared to the bulk soil [56]. Notably, only 48% of these global core taxa have genus classification, suggesting that most of them are under-described in their functional potential and interactions with plants (**Table S4**).

**Fig. 2:**
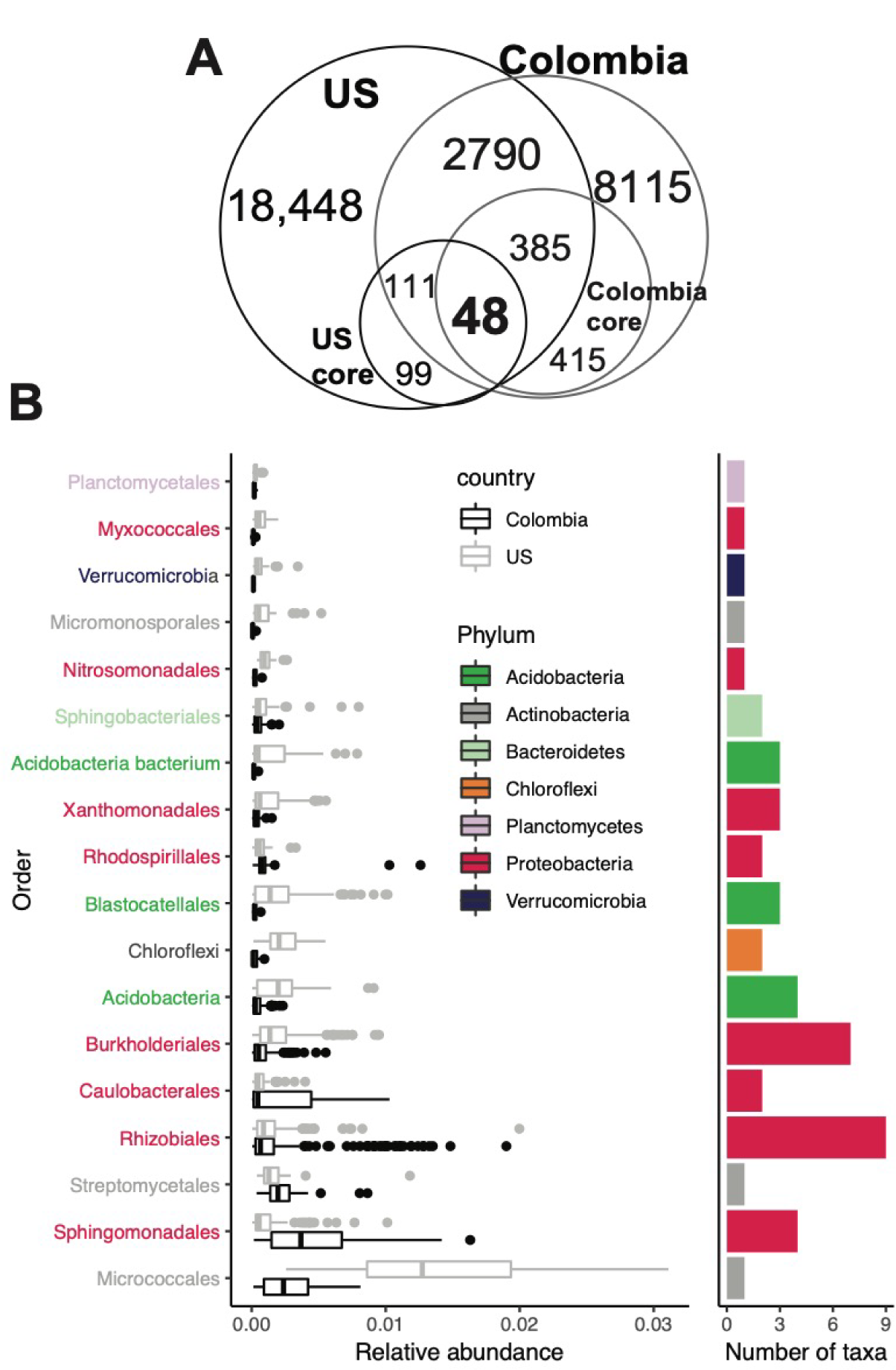
A global core rhizosphere microbiome. There were 3361 shared bacterial/archaeal taxa across U.S. and Colombia rhizosphere samples, suggesting highly similar recruitment across continental scales. Forty-eight taxa were detected in all samples of both datasets, as depicted by the Venn diagram (A), and included many Proteobacteria (B). Relative abundance of the 48 U.S.-Colombia core taxa is represented as boxplots (left panel), grouped by order and dataset (Colombia/U.S.). Number of taxa per order is represented as bars (right panel). Labels on the y-axis and bars are color-coded by phylum level.

We also analyzed reads associated with the global core taxa with the UNOISE3 pipeline [28] to generate predicted biological sequences (zero-radius OTUs – ZOTUs with100% sequence identity) and provide the maximal possible biological resolution. We found that the global core taxa consisted of 422 ZOTUs, and that there was a range of 2 to 35 ZOTUs identified within each OTU (**Fig. S5**). With one exception (HQ597858.1.1508), all of core OTUs (clustered at 97% sequence identity) contained at least one ZOTU that also had an occupancy of 1. In addition, all of the ZOTUs with an occupancy of 1 were also the most abundant ZOTUs within each OTU (**Fig. S5**). This result suggests that the same members constitute the core even with increased taxonomic resolution.

A recent study considered the effect of the common bean domestication history on the root microbiome and identified a core set of microbial taxa that are consistently present with these diverse bean genotypes, including bacterial taxa recruited from agricultural and natural soil from Colombia [15]. This core also had high representation of *Proteobacteria, Acidobacteria, Actinobacteria* and *Verrucomicrobia*, similar to those observed in the present study. We also re-analyzed these data and show that 46 (out of 48) of the global core taxa identified here have occupancy>0.9 across all three included studies (the present study, Pérez-Jaramillo et al. [14] and Pérez-Jaramillo et al. [15]). Forty-two of these taxa had the highest possible occupancy of 1, but only within rhizosphere samples from agricultural soils (**Table S4**). When incorporating data from beans grown in forest soils, the average occupancy decreases to 0.77 (median 0.75, min = 0.53). However, 14 taxa still had an occupancy of 1 when including beans grown in forest soils (**Table S4**).

In summary, these results show that common bean can associate with a core set of rhizosphere microbiome members at taxon and ecotype levels, across diverse bean genotypes and across continents. Additionally, these core taxa are likely enriched by the host in managed soils, as suggested by their higher occupancy in agroecosystems.

### Core taxa are enriched in the rhizoplane and are consistently detected across bean development

To identify the common bean core taxa over space (a biogeographic core) we sampled plants across growing locations at the same growth stage (flowering) and focused on the rhizosphere compartment. However, the question remained whether these core taxa are detected beyond that particular plant development stage. To answer this question, we conducted a field experiment to assess the core taxa over time in plant development stage. In the next growing season (2018), we used the same divergent bean genotypes grown at both Michigan, U.S, locations (Montcalm and Saginaw Valley; see Material and Methods). We harvested root systems at 5 different plant development stages including flowering stage. We investigated the relative abundance of the global core taxa and the U.S.-specific core taxa in both rhizosphere (soil that could be removed from the root after shaking, n=125 samples) and rhizoplane (soil adhered to the root tissue and removed via vortex in buffer, n=127 samples) compartments to determine their ability to closely associate with the plant tissue. The range of the rhizosphere sequencing depth was 7,905-78,436 reads per sample, and for the rhizoplane it was 32-189,433 reads per sample. We rarefied to 15,000 reads per samples (i.e. samples reaching richness asymptote), resulting in loss of 3 rhizosphere and 7 rhizoplane samples for a final dataset of 122 rhizosphere and 120 rhizoplane samples. The total richness observed was 36,022 bacterial and archaeal OTUs.

From this development time series, we found that all 48 global core taxa were detected a year later on these two Michigan farms. The collective relative abundances of the global core taxa were significantly higher in the rhizoplane as compared to the rhizosphere irrespective of the plant development stage, bean genotype and growing location (**Fig. 3A**). Interestingly, the remaining US core taxa that were found exclusively in the U.S. dataset at an occupancy of 1 were equally abundant in the rhizosphere and rhizoplane at both Michigan growing locations (**Fig. 3A**).

**Fig. 3:**
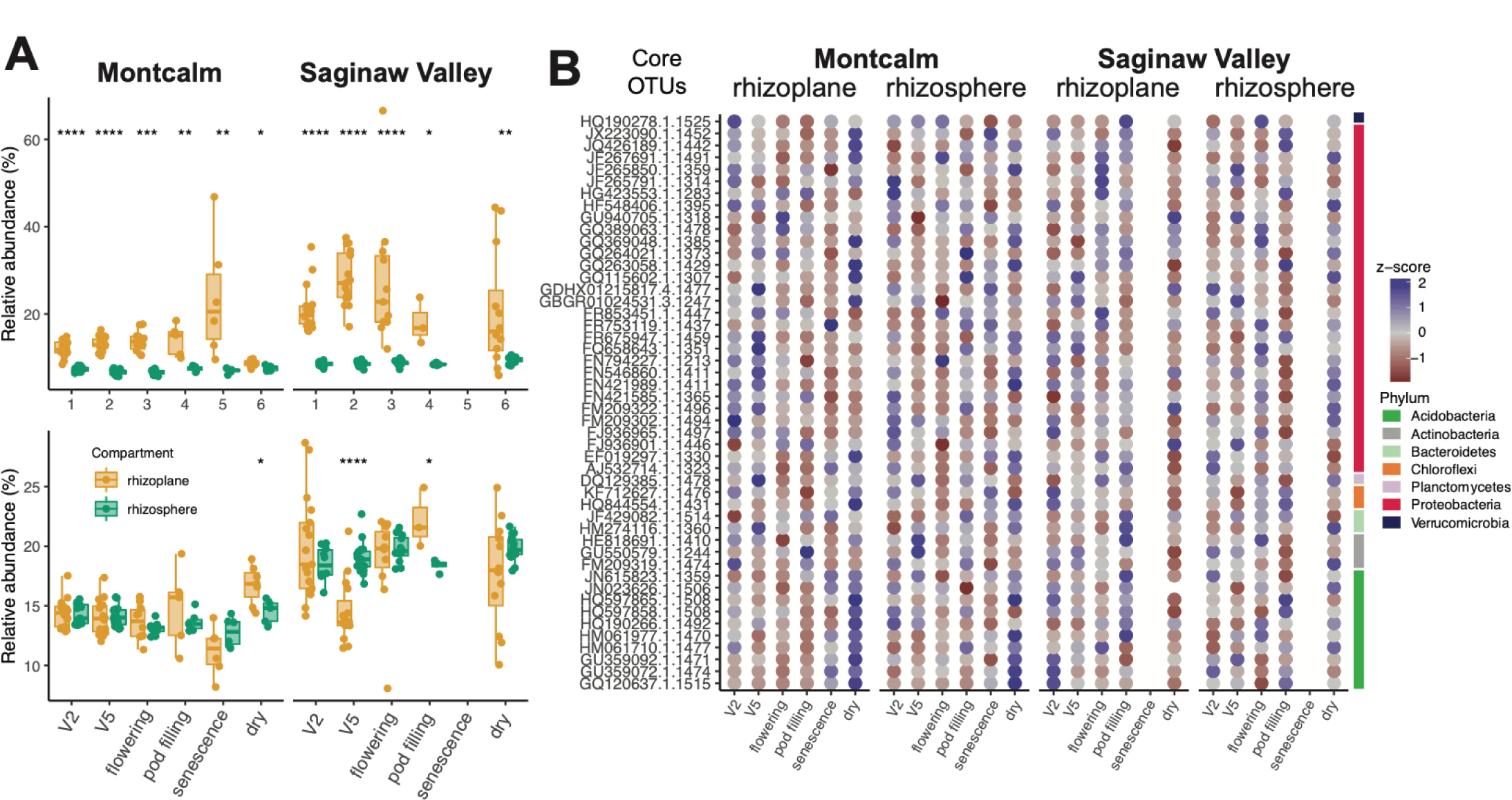
Relative abundance of core taxa in the root system of the common bean during plant development by root compartment and growing location. The combined relative abundance of 48 core taxa is significantly higher in the rhizoplane (green) compared to the rhizosphere (orange) and does not show plant development dependence (A – upper panel). The combined relative abundance of remaining core taxa specific to the U.S. (n=210), tend to be high throughout the plant development but are equally abundant in both compartments (A – lower panel). Stars above box plots represent statistical significances as determined by Wilcoxon test (**** ≤ 0.0001, *** ≤ 0.001, ** ≤ 0.01, * ≤ 0.05). Z-score normalized relative abundance of the 48 core taxa across plant development, compartments and growing location (B). Taxa on the y-axis are arranged by their classification at the phylum level.

We next asked whether there were enrichments of particular core taxa by plant development stage, root compartment, or growing location (**Fig. 3B**). On balance, almost all core taxa showed some growth stage preference, but these trends were specific to each growing location and root compartment. Despite these nuances, all core taxa were consistently found with high occupancy inclusive of the plant development series (**Fig. S6**).

Together these results suggest that these core taxa are selected by the plant early in the development stage and maintained. Enrichment of the core taxa in the rhizoplane further supports the hypothesis that these core taxa engage closely with the host plant.

### Core taxa are not hub or connector taxa in an inter-domain microbiome network

Network analysis has been proposed to be a useful method to identify important members of the plant microbiome with beneficial traits [57–59]. Hub taxa, identified by their high connectivity with many members of the community, are regarded as the most important part of the community and influence network structure and community stability [60, 61]. We applied this method to ask if any of the core taxa that we identified using abundance-occupancy were also key for co-occurrence network structure. Additionally, we were interested in identifying fungal-bacterial co-occurrences because of reports of their potential benefits for the plant [62, 63]. To explore these patterns we applied the molecular ecology network analysis pipeline (MENAP) which constructs ecological association network through random matrix theory (RMT) [30]. We merged rarified 16S and ITS rhizosphere datasets, filtered the datasets to include taxa with occupancy greater than or equal to 50%, and considered only interactions significant at *p*-value<0.05 and RMT threshold of 0.88. The resulting network included 572 taxa (nodes) and 1,857 statistically significant co-occurrences (edges) structured among 52 modules. Most of the modules were relatively small, with only six including more than 10 nodes (**Fig. 4**). The network was scale-free (i.e. characteristics of the network are independent of the size of the network) and had small-world characteristics (i.e. highly clustered) as indicated by the node degree distribution fitting to the power law model (R^2^= 0.993), and also had significant deviation of the modularity, length and clustering coefficients from those calculated from random network (i.e. same number of nodes and edges), respectively (**Table S5**).

**Fig. 4:**
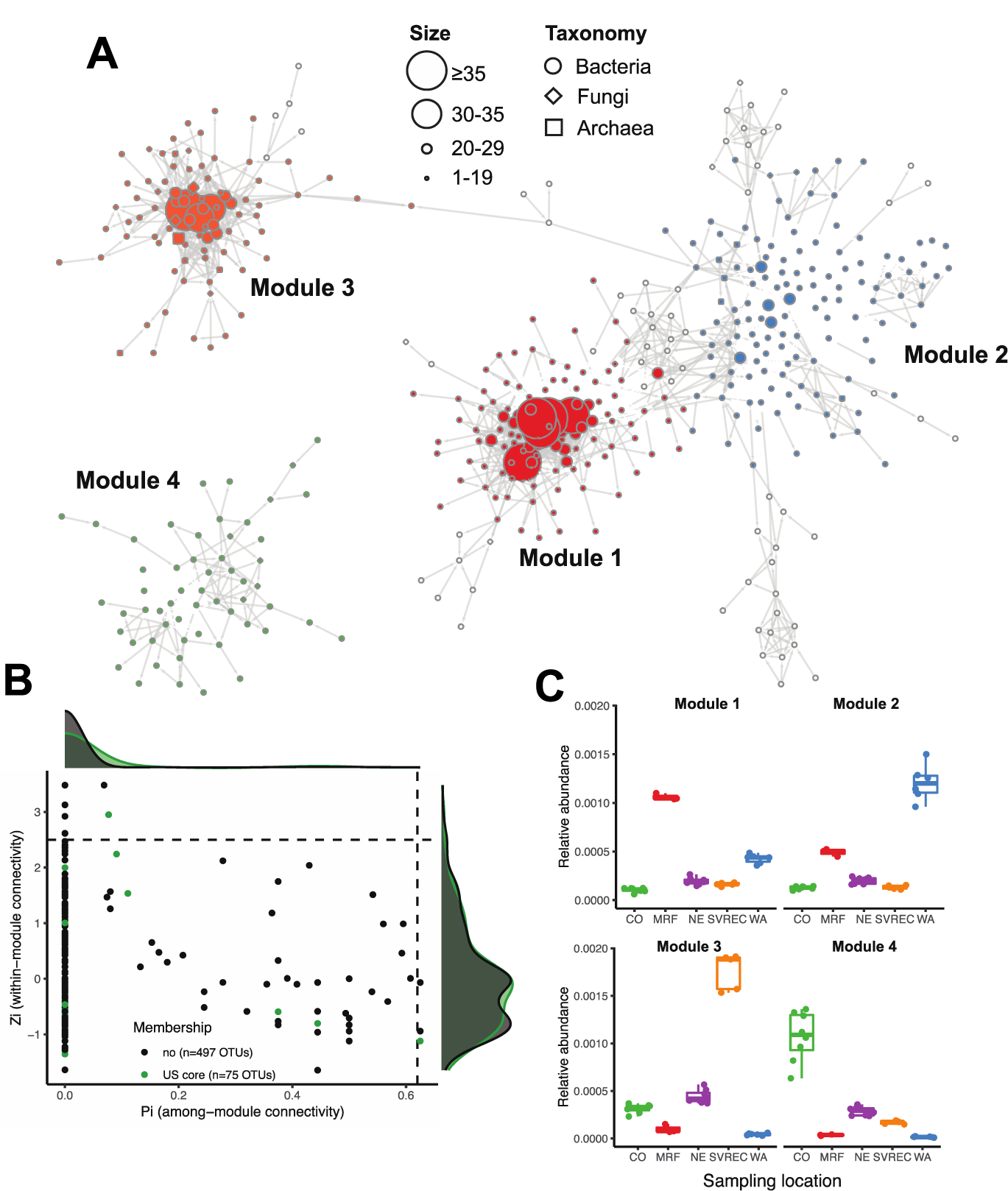
Network co-occurrence analysis shows that core rhizosphere microbiome members are predominantly classified as peripheral taxa that are weakly connected, and generally clustering by the growing location. The network depicts only clusters of modules that were connected by more than 6 nodes (A). Nodes shape is representing domain associations (archaeal, bacterial or fungal) and node size is proportional to its total number of connecting edges (A). The four largest modules are generally reflective of community biogeography and distinguished by color (A). The within-(Zi) and among-(Pi) module connectivity plot was used to identify module (Pi<0.62, Zi>0.2.5) or network hub taxa (Pi>0.62, Zi>2.5), as well as connector (Pi>0.62, Zi<2.5) and peripheral taxa (B). The density plots surrounding the Zi-Pi plot represent core (green) and non-core taxa (black) (B). The relative abundances of taxa within each module is represented in box plots (C).

The topological role of each taxon within the network was determined by the relationship between their within-module (Zi) and among-module (Pi) connectivity scores as described in [64]. Based on this, the majority of taxa were peripheral (potentially, specialists; 563 nodes) (**Fig. 4A**). There were also 3 connectors and 6 module hubs, but no network hubs. Indeed, in agreement with the beta-diversity analysis (**Fig. S3**, Supporting Materials), there was a strong geographic signal in the largest four modules, and these were comprised mostly of bacterial-bacterial (rather than bacterial-fungal or fungal-fungal) hypothesized interactions (**Fig. 4B**). We note that cross-domain edges constituted only a small fraction of all co-occurrences (n=168; bacteria-fungi=156, archaea-bacteria=6 and archaea-fungi=6).

The analysis identified 26 co-occurrences between core taxa. However, we were surprised to find that only two bacterial core and no fungal core taxa were also classified as network hubs (taxa that connect to many other taxa within a modules) or connectors (taxa that connect across modules). As exceptions, core *Chitinophagaceae* taxon FR749720.1.1492 was a module hub node and a *Nitrobacter* sp. GDHX01215817.4.1477 was a connector. Our results, inclusive of a dataset of divergent plant genotypes and broad biogeography, suggest that while hub and connector taxa may be important for the maintenance of the root microbiome, these taxa are not consistently detected in the common bean rhizosphere and, by deduction, could not be of universal importance for the host plant. Our study cannot speak to the potential for functional redundancy among hub or connector taxa, which could ultimately suggest a functional core among phylogenetically diverse taxa [65]. Taken together, these results suggest that core taxa likely are important for the plant, while hub and connector taxa are important for the integrity of the soil microbial community and its responses to the local environment.

## Discussion

Numerous studies have aimed to detect core microbiome members for various plants and animals (see references within [29, 66]). Because methods are inconsistent across studies and because the parameters used to define a core are often arbitrary, there is some question as to how robust or useful such studies may be. For these reasons, it has become easy to overlook new studies that claim to have discovered a core microbiome for their system. If every study or design results in a different core microbiome, how can the research build to move forward?

Here, we applied concepts from macroecology to prioritize, for the first time, a core plant microbiome that persists at across growing locations on two continents, over plant development, over two years of annual plantings at two different farms, across highly divergent plant genotypes, and also across datasets collected by different research groups. Furthermore, core members were enriched in the rhizoplane, and not just the rhizosphere. These multiple and consistent lines of evidence provide strong support that these taxa likely engage with the plant and also demonstrate the robustness of the abundance-occupancy method to discover core members.

A majority of the 48 core bacterial taxa among these common bean rhizospheres are under-described or largely unknown, as only 23 out of 48 have a genus-level classification. But, due to their ubiquitous association with the bean over space and time, we hypothesize that these taxa provide functions that are crucial for common bean health and should be targeted for microbiome management of in support of common bean productivity and wellness. To test this hypothesis further research is needed, but plant-beneficial traits have been previously reported for members of the genera that we identified as part of the bean core microbiome. For example, members belonging to the genera *Mesorhizobium* and *Rhizobium* are known symbiotic nitrogen fixers of legumes [67, 68], *Ramlibacter* sp. have the ability to promote P mobilization [69], and *Variibacter, Novosphingobium* and *Sphingomonas* sp. harbor many specialized genes indicating their relationship with plant hosts [70–72].

Discovering these core taxa is a first step in a rich line of inquiry to understand host engagement with them. The next steps are to understand functions associated with these taxa and to determine how they contribute to plant health and productivity under different growth conditions, such as drought or with particular management strategies [39, 73]. These steps will include cultivation dependent and independent approaches aimed to enrich and isolate core members, assemble or bin genomes from isolates and metagenomes, annotate functional genes on both chromosomes and plasmids, link functions and activities through transcript or metabolome analyses, and perform experiments with constructed communities of core members to test hypotheses about microbiome engagement with and benefits to the plant.

To conclude, this work provides robust approaches and general insights for prioritizing core microbiome members, and that also advance goals in plant-microbiome management and microbe-improved crops by providing insights into core member identities and dynamics.

## Supporting information

Supporting Methods, Results, and Discussion

## Acknowledgments

This work was supported by the Plant Resilience Institute at Michigan State University. We are grateful to Dr. James Kelly, Dr. Phillip Miklas, Dr. James J. Heitholt, Dr. Carlos Urrea Florez, Dr. Mark A. Brick, Dr. Juan M. Osorno, Dr. Thomas H Smith and other supporting staff at the research extension farms for their partnership in this study. We would also like to thank the Michigan State Genomics Core Research and Technology Support Facility and the Institute for Cyber-Enabled Research High Performance Computing Center for excellent support and service. AS acknowledge support from Michigan State AgBioResearch (Hatch).

## Author contributions

N.S. and A.S. designed research, analyzed data and wrote the paper; and N.S. performed research.

## Supporting Figure legends

**Fig. S1**: Sequencing depth and rarefaction curves for 16S rRNA (A,B) and ITS (C, D) dataset from samples collected in 2017. The red splitted line represents the rarefaction threshold. Note that we submitted amplicons for sequencing from the 2017 sampling effort twice which resulted in the sample to sample variation in read depth (rhizosphere 16S rRNA samples sequenced first followed by the ITS amplicons with addition of the root-associated samples).

**Fig. S2**: Alpha diversity indices for the 16S RNA (A, B) and ITS (C, D) datasets. Represented are rich-ness, Shannon and Pielou indices measured by growing location (A, C) and bean genotype (B, D). For statistical comparison of the pairs we used ANOVA (A, C) or Wilcoxon test (B, D).

**Fig. S3**: Growing location drives bacterial/archaeal (A) and fungal (B) microbiome structure of the common bean rhizosphere. The principal coordinate analysis (PCoA) is based on Bray-Curtis distances. Growing location is indicated by color and plant genotype is indicated by shape shapes (diamond=CELRK, circle=Eclipse, square=root zone soil). The strength of statistically significant (p-value < 0.01) explanatory variables are shown as the length of fitted vectors.

**Fig S4**: Community composition (A, C) and number of shared taxa between sites represent as Venn diagrams (B, D). The 16S rRNA dataset is represented in the top panels (A, B) and ITS in the lower panels (C, D). The bar charts are colored based on the phylum association (phyla represented by relative abundance < .05 are grouped and labelled as other). For Venn diagrams, samples were grouped by the growing location and root zone samples were removed.

**Fig. S5**: Analysis of ZOTUs represented by each identified core OTU. 48 core OTUs were represented by as few as 4 ZOTUs and by up to 35 ZOTUs. For every OTU we found at least one ZOTU with occupancy = 1 and all of them, except of 2 ZOTUs, had also the highest relative abundance among them. Points are color coded by their presence, red representing those with occupancy < 1 and blue for ZOTUs with occupancy of 1. The OTUs on the x-axis are ordered alphabetically.

**Fig. S6**: Occupancy of core OTUs in the development study. Occupancy is represented by color and size.

**Fig. S7**: Comparison of relative abundance of the 48 core OTUs between the DNA isolation methods used in the development study (G=Griffith, P=PowerSoil). Statistical difference, determined by Wilcoxon test, is represented as star symbol (*<0.05, **<0.01, ***<0.001). The pints are color coded by sample they derived from.

**Fig. S8**: The effect of isolation method on alpha and beta diversity. For alpha diversity richness, Shannon and Pielou indices are presented (A). For the principal coordinates analysis, Bray-Curtis distance matrix was used. Symbols are colored by samples. Wilcoxon test was used to determine statistical differences between isolation methods for the alpha diversity metrices (*<0.05). PERMANOVA was used to determine the effect of isolation method on community structure.

## Supporting Table legends

**Table S1:** PERMANOVA results for the 16S rRNA and ITS data. Highly correlated or/and statistically significant values are highlighted in bold.

**Table S2**: 20 differentially abundant OTUs between the two plant genotypes as identified by using DESeq2 (Love et al. 2014).

**Table S3**: Sloan neutral model summary.

**Table S4**: List of 48 core taxa and their taxonomic classification.

**Table S5**: Occupancy of core OTUs in agricultural, natural (forest) soils or when combined. Results are based on the re-analysis of the data from the Pérez-Jaramillo et al. 2019. OTUs with occupancy of 1 in both soils are highlighted in orange. Additional to the number we used green gradient shading to represents the occupancy of each OTU.

**Table S6**: Summary of network properties of actual and random network.

