## Supporting Methods, Results, and Discussion for "Prioritizing persistent microbiome members in the common bean rhizosphere: an integrated analysis of space, time, and plant genotype"

Supporting Results and Discussion

*Beta diversity: strong biogeographic but weak plant genotype influence on microbiome dynamics*

As expected, there were differences across growing locations in edaphic soil properties (e.g. pH, nitrogen species, organic matter) as well as in management practices (e.g. fertilization, irrigation, crop rotation) and climate (**Table 1**). Growing location, pH and fertilization explained differences in microbial community structure for bacteria/archaea and fungi, but plant genotype did not (**Fig. S3AB**, **Table S1**). Fungal and bacterial/archaeal communities had synchronous biogeographic patterns (Procrustes m12 squared: 0.2137, R^2^=0.8867 and *p*-value=0.001), supporting that both communities are shaped by local edaphic factors. An effect of growing location on beta-diversity has been previously reported for plant- and soil- associated microbiomes (1–4).

We selected two distinct bean genotypes with divergent evolutionary and breeding histories to assess the impact of plant genotype on the root microbiota, expecting that any signal would be maximized between these lineages. There was a weak interaction between plant genotype and growing location in explaining fungal community structure (PERMANOVA, R^2^=0.0377, *p*-value=0.001) (**Table S1**), but no effect of plant genotype alone on the bacterial and archaeal communities, even when controlling for growing location. Similarly, plant genotype had a weak but statistically detectable effect on bacterial and archaeal richness (*p*-value=0.019) but no measurable effect on fungal richness (*p*-value=0.95) (**Fig. S2BD**). An analysis of differential taxon abundances across genotypes (DESeq2 (5)) detected 20 taxa out of 21,881 total that were distinguishing between the genotypes CELRK and Eclipse, but their adjusted *p*-value were marginally significant (*p*-value> 0.02; **Table S2**). The absence of a robust genotype effect on the microbiome is in contrast to a previous study on common bean (6). However, several studies have reported a relatively weak to no influence of genotype more generally on the rhizosphere communities of other plants grown in the field (1, 2, 7). The study that observed a genotype effect on the bean root microbiota included wild, domesticated and landrace common beans of Mesoamerican origin that were grown in a greenhouse using Colombian field soil (6). The authors attributed those bean genotype differences to differences in root architecture, which often has a different phenotype in greenhouse pots than in the field (8). We summarize that, in this study and in agreement with other field studies, plant genotype has a minor to no measurable impact on rhizosphere microbiome.

Supporting Materials and Methods

*Soil chemical analysis*

Soil analysis was done at the Michigan State Soil and Plant Nutrient Laboratory by agricultural soil testing standard protocols and included following parameters: pH, phosphorous (P ppm), potassium (K ppm), calcium (Ca ppm), magnesium (Mg ppm), organic matter content (OM%), nitrate (NO_3_^-^ ppm), ammonium (NH_4_^+^ ppm) and total nitrogen (%).

*DNA isolation and sequencing*

For the biogeography samples collected in 2017, DNA was isolated using DNeasy PowerSoil kit (QIAGEN, US) by following the manufacturer’s recommendations. For the plant development samples collected in 2018, DNA was isolated using Griffiths protocol (9) except for the SVERC rhizosphere for which DNeasy PowerSoil kit (QIAGEN, US) was used because the Griffith protocol had low nucleic acid yield. Thus, we also assessed the effect of DNA isolation protocol on amplicon sequencing outcomes by comparing eight MRF rhizosphere samples for which DNA was isolated using both methods. Analyses suggest that the relative abundance of only 12 out of 48 core taxa was significantly influenced by the isolation method (**Fig. S7**) thus we believe that the comparison of relative abundance of core taxa, between the root compartments in SVERC is possible. Notably, isolation methods influenced both alpha (richness but not Shannon and Pielou) and beta diversity (**Fig. S8**).

Quality and quantity of isolated DNA was assessed with Qubit 2.0 fluorometer using HS dsDNA Assay kit (ThermoFisher, US). Presence of 16S rRNA genes was confirmed by PCR using the V4 16S rRNA sequencing primer set 515f and 806r (10) and then visualizing the PCR products by agarose gel electrophoresis. In 2017 for the spatial study, rhizosphere samples were pooled per location by field plot. In 2018 for the temporal study, rhizoplane and rhizosphere samples were assessed for each individual plant harvested. 16S rRNA gene amplicon samples were prepared by the Michigan State Genomics Core Research Support Facility. Their standard protocol included 16S rRNA gene amplicon PCR amplification and library preparation. The ITS samples were amplified in our lab using the primer pair ITS1f and ITS2 (11) with index adapters as recommended Genomics Core (https://rtsf.natsci.msu.edu/genomics/sample-requirements/illumina-sequencing-sample-requirements/, June 2019), The Genomics Core used the Illumina TruSeq Nano DNA library preparation kit for both 16S and ITS libraries. Paired-end, 250-bp reads were generated on an Illumina MiSeq platform using v2 Standard 500 cycle kit, and the Genomics Core provided standard Illumina quality control, adaptor, barcode trimming, and sample demultiplexing with Illumina Bcl2fastq v2.19.1. In all sequencing efforts we included blank sample (negative control) which were processed the same way to identify OTUs resulting from contamination either through DNA isolation or PCR.

*Statistical analysis*

Alpha and beta diversity analyses were performed to datasets subsampled to the minimum observed quality filtered reads per sample (2017 dataset: 31,255 for 16S rRNA and 22,716 for ITS, 2018 dataset: 15,000 for 16S rRNA). Due to low read counts one sample (SVERC1) was removed from the 16S rRNA 2017 dataset. We report richness as total number of OTUs clustered at 97% sequence identity. Differences in alpha diversity among groups (plant genotype, root compartment, sampling time, growing location, fertilization, pH) were assessed using analysis of variance (ANOVA), with a Tukey *post hoc* test for multiple comparisons. We used the *protest* function in the vegan package in R (19) (version 2.5-6) to test for synchrony between bacterial/archaeal and fungal communities.

To quantify the variation in beta diversity (community structure), we first calculated pairwise Bray-Curtis dissimilarities. Permutational multivariate analysis (PERMANOVA) using 1,000 random permutations was used to test hypothesis of beta diversity using *adonis* function in the vegan package in R (19).

We used DESeq2 (5) to determine bacterial and archaeal taxa with differential abundance across plant genotypes from the biogeography study. Thresholds for calling taxa as differentially abundant between selected groups was set at p-value of 0.05.

We prioritized core taxa over space and time using abundance-occupancy distributions fitted to the Sloan neutral model, as recently described (20). Species abundance-occupancy distributions are often applied to explore large-scale patterns in species distributions, especially in macroecology (21). For that, first taxon’s mean relative abundance is calculated across the dataset and log transformed and secondly, its frequency of detection across dataset is calculated (termed as occupancy, with 1 represented in all samples). In this study we considered as core all taxa that were found in every sample across growing locations. To assess the importance of neutral process in the assembly of common bean rhizosphere communities we applied the Sloan neutral model (22). This model predicts the relationship between the frequency with which taxa occur in a set of local communities (occupancy) and their mean abundance across a broader metacommunity. The prediction is that rare taxa will be lost from individual hosts due to ecological drift, but abundant taxa will be more widespread in a metacommunity due to higher chance of dispersal and thus be randomly sampled by individual hosts. The occupancy of OTUs and their mean relative abundances across the metacommunity were fitted to the model, using the R code described by Burns et al. (23). We used the following parameters of neutral model: 95% confidence intervals, the goodness of fit of the neutral model (R^2^), and the estimated migration rate (m).

*Co-occurrence network analysis*

Global network properties were calculated using the Molecular Ecological Network Analysis Pipeline (MENAP) (24). The majority was set to 0.5, missing data were kept blank, read counts were converted by the logarithm, and the Pearson correlation coefficient was used as the similarity measure. The results were then filtered by keeping pairwise correlations with absolute LA values greater or equal to 0.88. To determine the modularity of the network we created 100 random networks within MENAP using the same number of nodes and edges as the complete potentially active network. The general properties of the inferred networks were analyzed using NetworkAnalyzer in Cytoscape (25, 26).

The connectivity of each node in the network was calculated using within-module connectivity (Zi) and among-module connectivity (Pi) scores which defined the topological role of each node (taxon) (24, 27). We classified our nodes as per the four categories describing node topology: *network hubs* (highly connected nodes within the entire network, Zi > 2.5 and Pi > 0.62), *module hubs* (highly connected nodes within modules, Zi > 2.5), *connectors* (nodes that connect modules, Pi > 0.62), and *peripheral* nodes (nodes connected in modules with few outside connections, Zi < 2.5 and Pi < 0.62) (27).

Cytoscape v.3.5.1 (25) was used for visualization of significant co-occurrences and editing the appearance of nodes size, shape and color based on the number of connections (degrees), taxonomic affiliation and module, respectively.

*Comparative analyses with published datasets*

To determine whether members of the US bacterial and archaeal core were associated more generally with bean plants grown, we compared our data to a published study of bean rhizosphere conducted in Colombia that investigated the influence of common plant genotype and root architecture on the rhizosphere microbiome composition. This published study included eight Mesoamerican genotypes planted in an agricultural soil (Colombia forest), and grew the plants in the greenhouse (6) (NCBI BioProject ID PRJEB19467), We downloaded and processed raw reads as described above, and, despite differences in our processing pipeline and the originally published pipeline, we generated very similar numbers of OTUs (12,209 compared to 12,293 (6)). Because the studies used different sequencing primers, we identified identical OTUs by matching the IDs provided by the SILVA database v128 (15) for all reference clustered OTUs. For the *de-novo* clustered OTUs, we used BLAST (28) to identify 100% matches between the two datasets, across the overlapping V4 region of reads from both studies. As before, we calculated occupancy of every taxon in the rhizosphere datasets and identified those with occupancy of 1. Taxa with the same OTU IDs or 100% BLAST match were designated as the cross-continental core rhizosphere microbiota of the common bean.

A similar approach was used to compare the data published in Pérez-Jaramillo et al. (29) (BioProject ID PRJEB26084). We combined this dataset with the one from above (PRJEB19467 (29)) and re-run the same UPARSE pipeline (version 11) (12) and classify the OTUs with SILVA database v128 (15). We attempted to re-analyzed these data as closely as to what was published by the authors, however, due to incomplete methodology description and metadata, we were not able to do so. Thus, we applied these published data to search for the core taxa occupancy in agricultural and forest soils.
