## Supplemental Figures for "Prioritizing persistent microbiome members in the common bean rhizosphere: an integrated analysis of space, time, and plant genotype"

FigS1

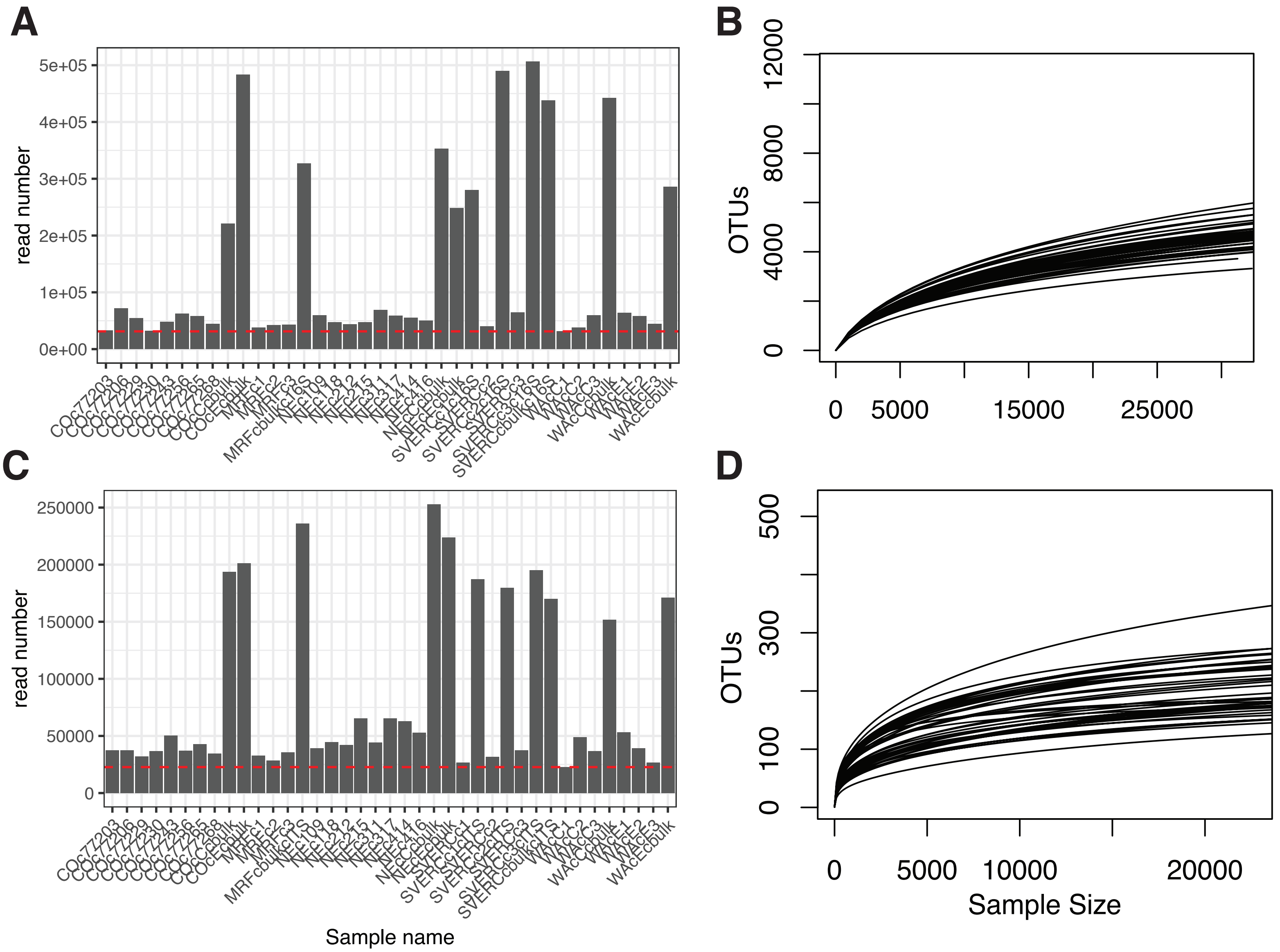


**Fig. S1**: Sequencing depth and rarefaction curves for 16S rRNA (A ,B) and ITS (C, D) dataset from samples collected in 2017. The red splitted line represents the rarefaction threshold. Note that we submitted amplicons for sequencing from the 2017 sampling effort twice which resulted in the sample to sample variation in read depth (rhizosphere 16S rRNA samples sequenced first followed by the ITS amplicons with addition of the root-associated samples).

FigS2

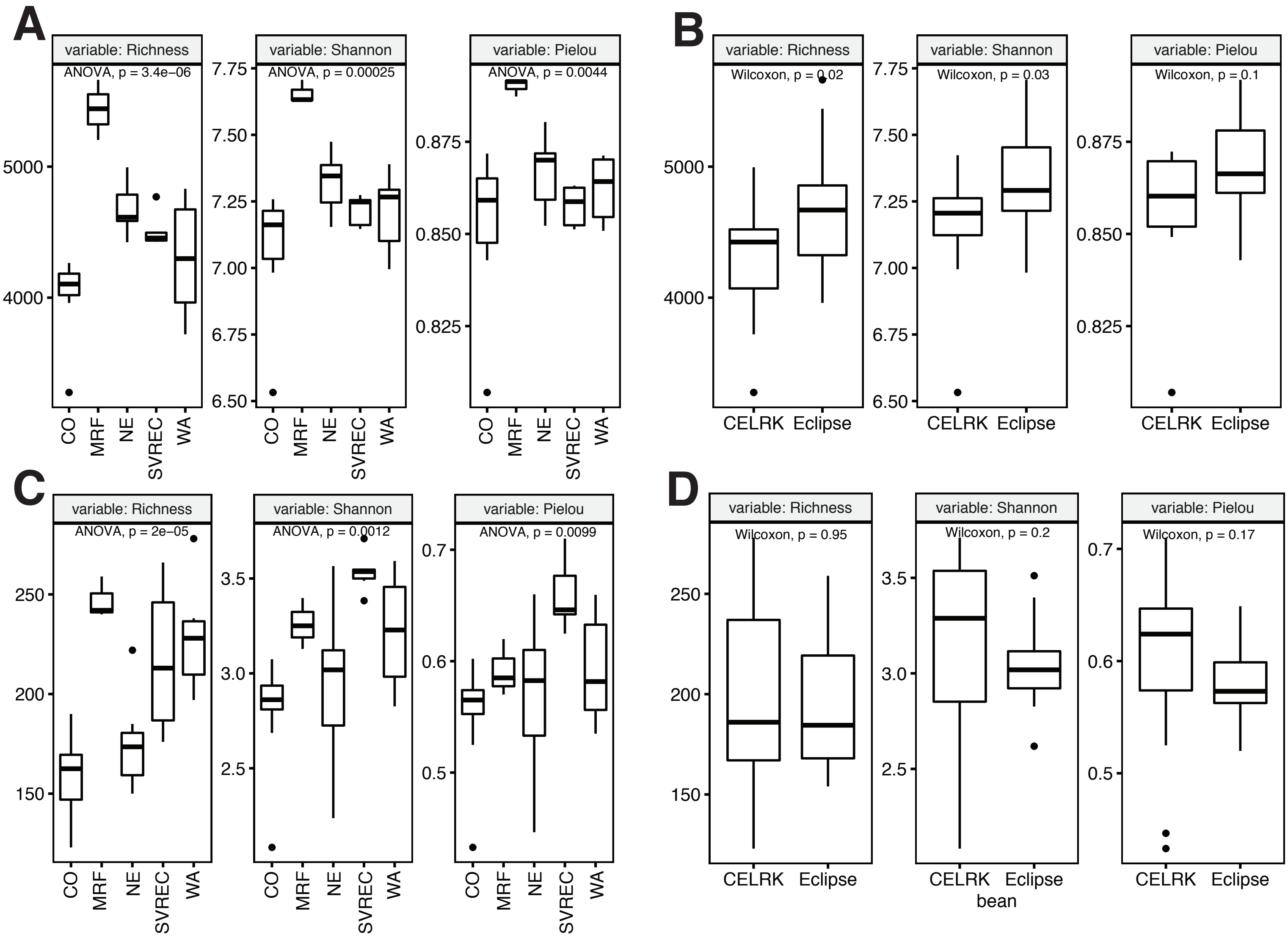


**Fig. S2**: Alpha diversity indices for the 16S RNA (A, B) and ITS (C, D) datasets. Represented are richness, Shannon and Pielou indices measured by growing location (A, C) and bean genotype (B, D). For statistical comparison of the pairs we used ANOVA (A, C) or Wilcoxon test (B, D).

FigS3

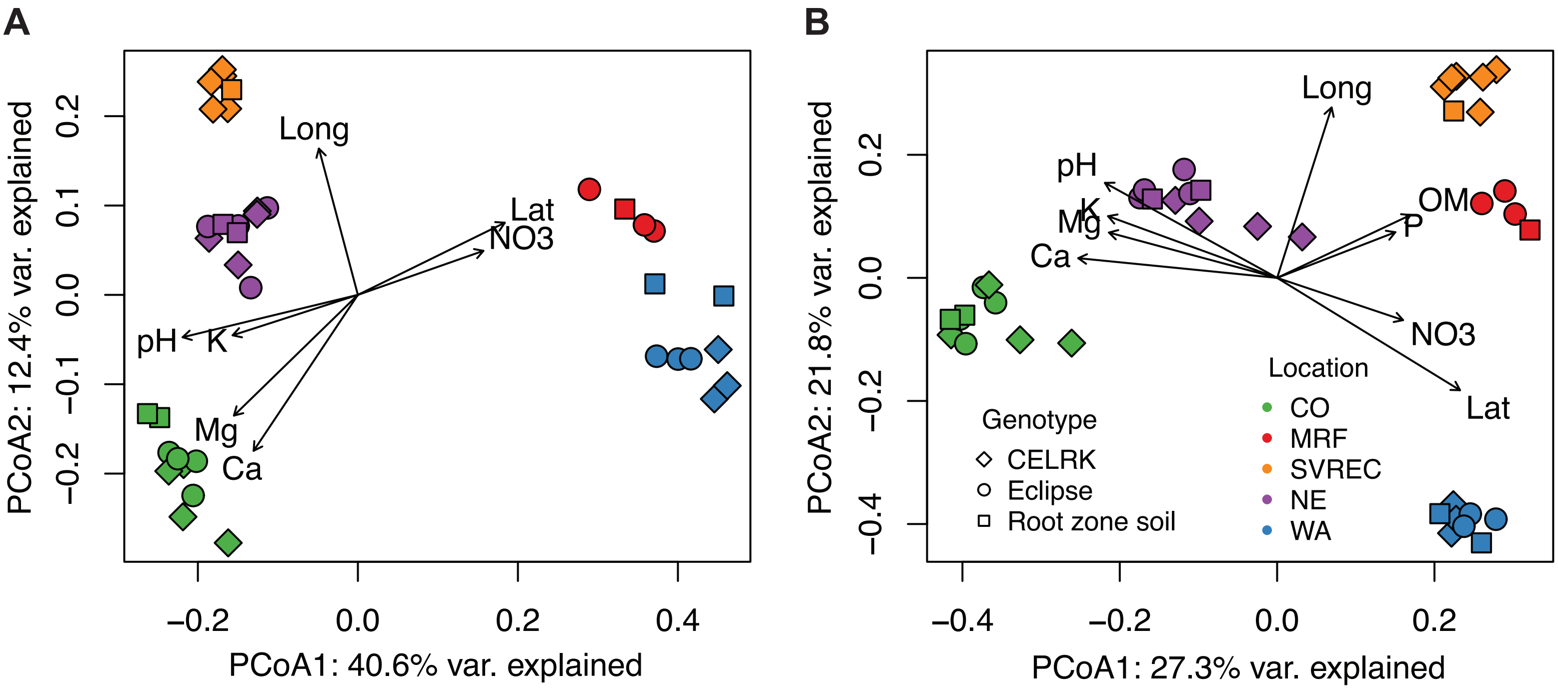


**Fig. S3**: Growing location drives bacterial/archaeal (A) and fungal (B) microbiome structure of the common bean rhizosphere. The principal coordinate analysis (PCoA) is based on Bray-Curtis distances. Growing location is indicated by color and plant genotype is indicated by shape shapes (diamond=CELRK, circle=Eclipse, square=root zone soil). The strength of statistically significant (p-value < 0.01) explanatory variables are shown as the length of fitted vectors.

FigS4

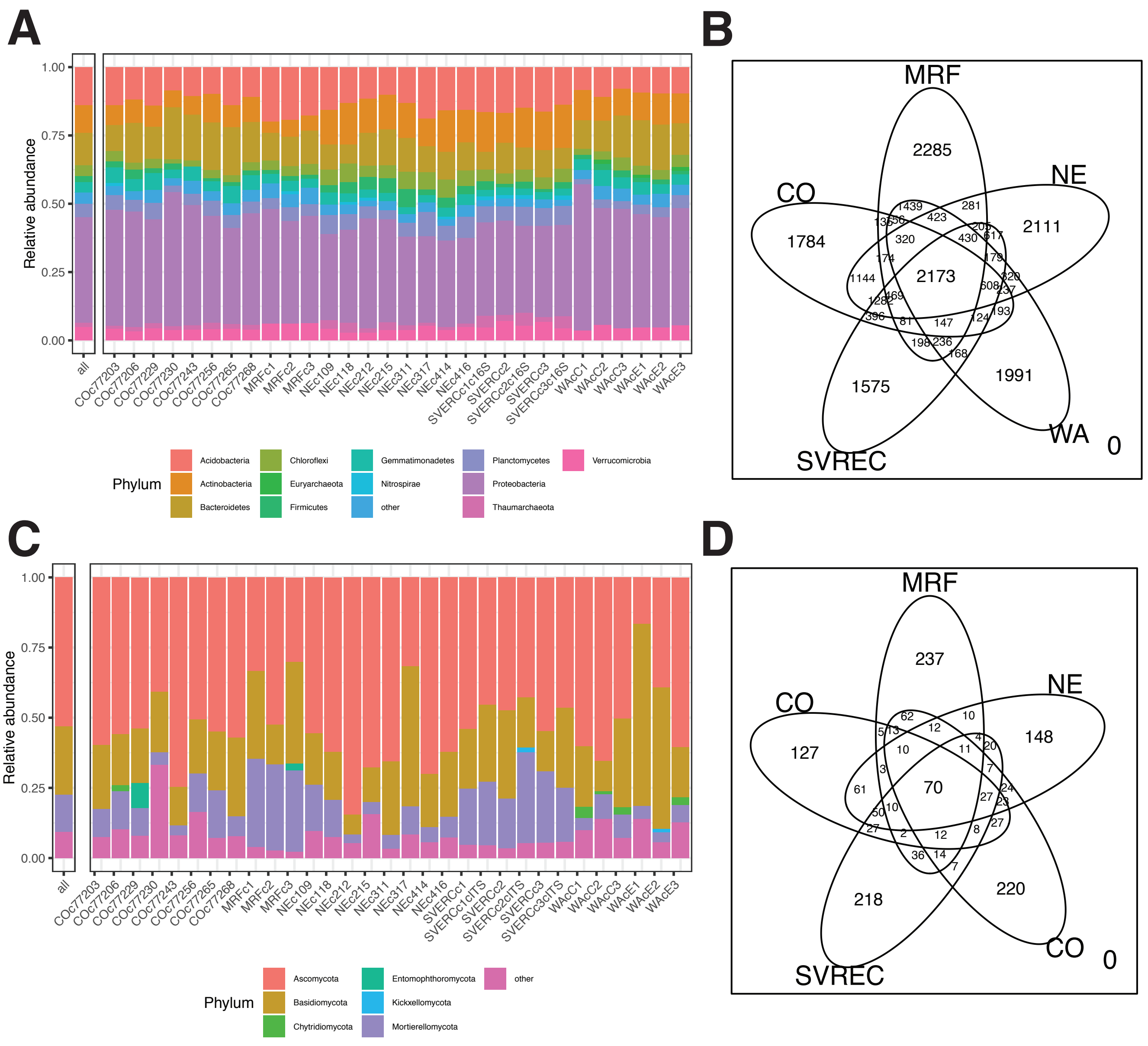


**Fig. S4**: Community composition (A, C) and number of shared taxa between sites represent as Venn diagrams (B, D). The 16S rRNA dataset is represented in the top panels (A, B) and ITS in the lower panels (C, D). The bar charts are colored based on the phylum association (phyla represented by relative abundance < .05 are grouped and labelled as other). For Venn diagrams, samples were grouped by the growing location and root zone samples were removed.

FigS5

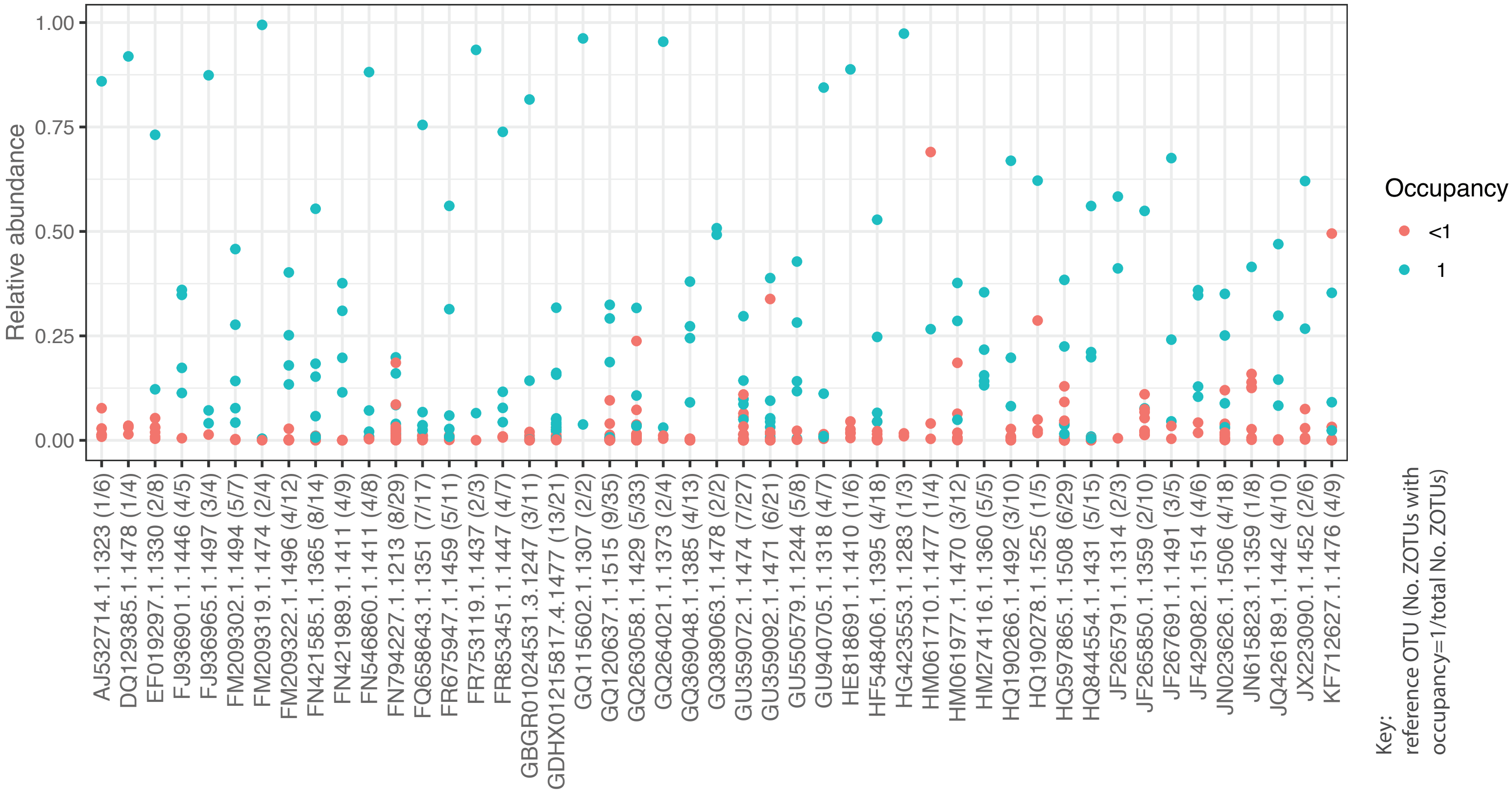


**Fig. S5**: Analysis of ZOTUs represented by each identified core OTU. 48 core OTUs were represented by as few as 4 ZOTUs and by up to 35 ZOTUs. For every OTU we found at least one ZOTU with occupancy = 1 and all of them, except of 2 ZOTUs, had also the highest relative abundance among them. Points are color coded by their presence, red representing those with occupancy < 1 and blue for ZOTUs with occupancy of 1. The OTUs on the x-axis are ordered alphabetically.

FigS6

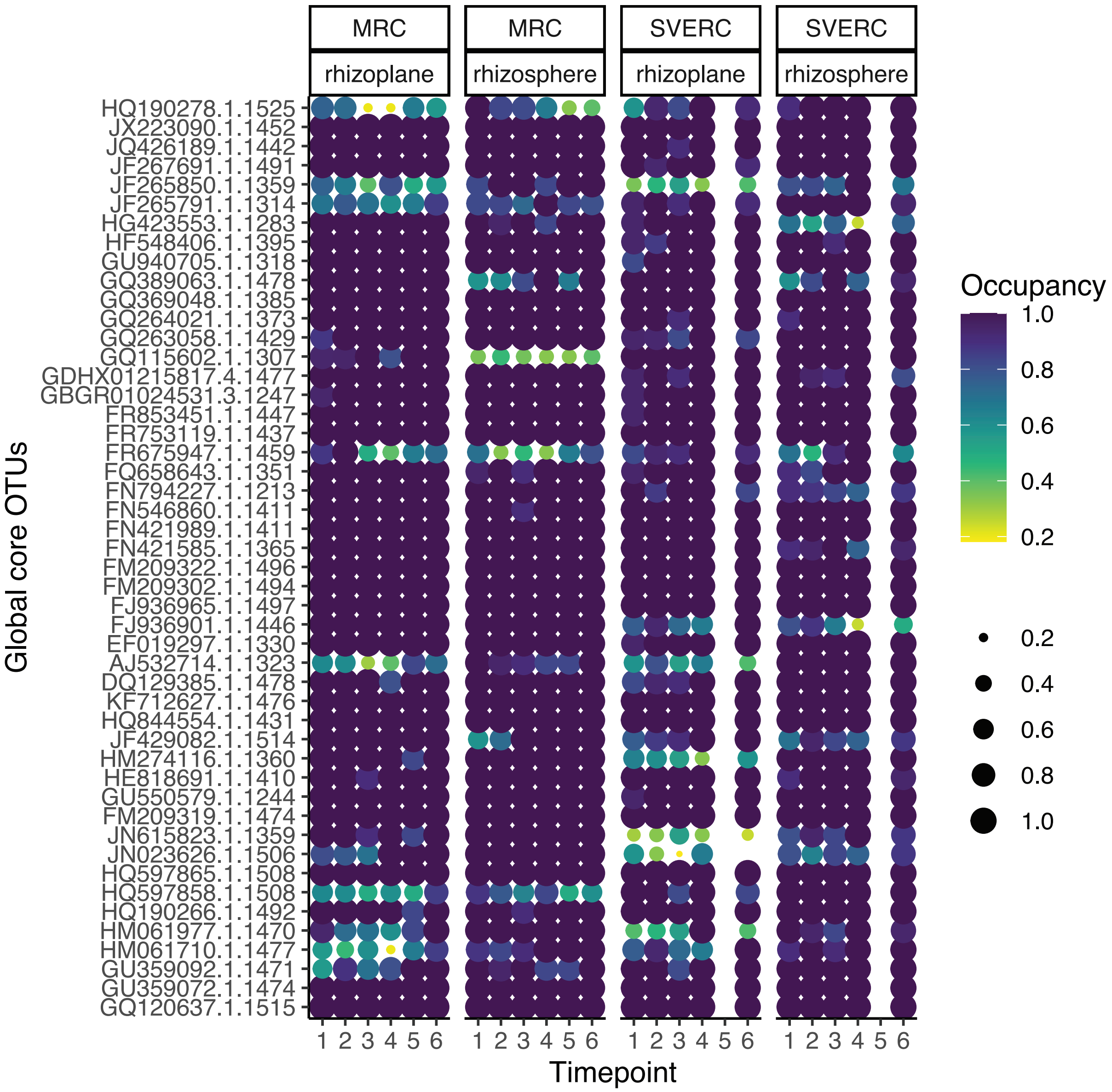


**Fig. S6**: Occupancy of core OTUs in the development study. Occupancy is represented by color and size.

FigS7

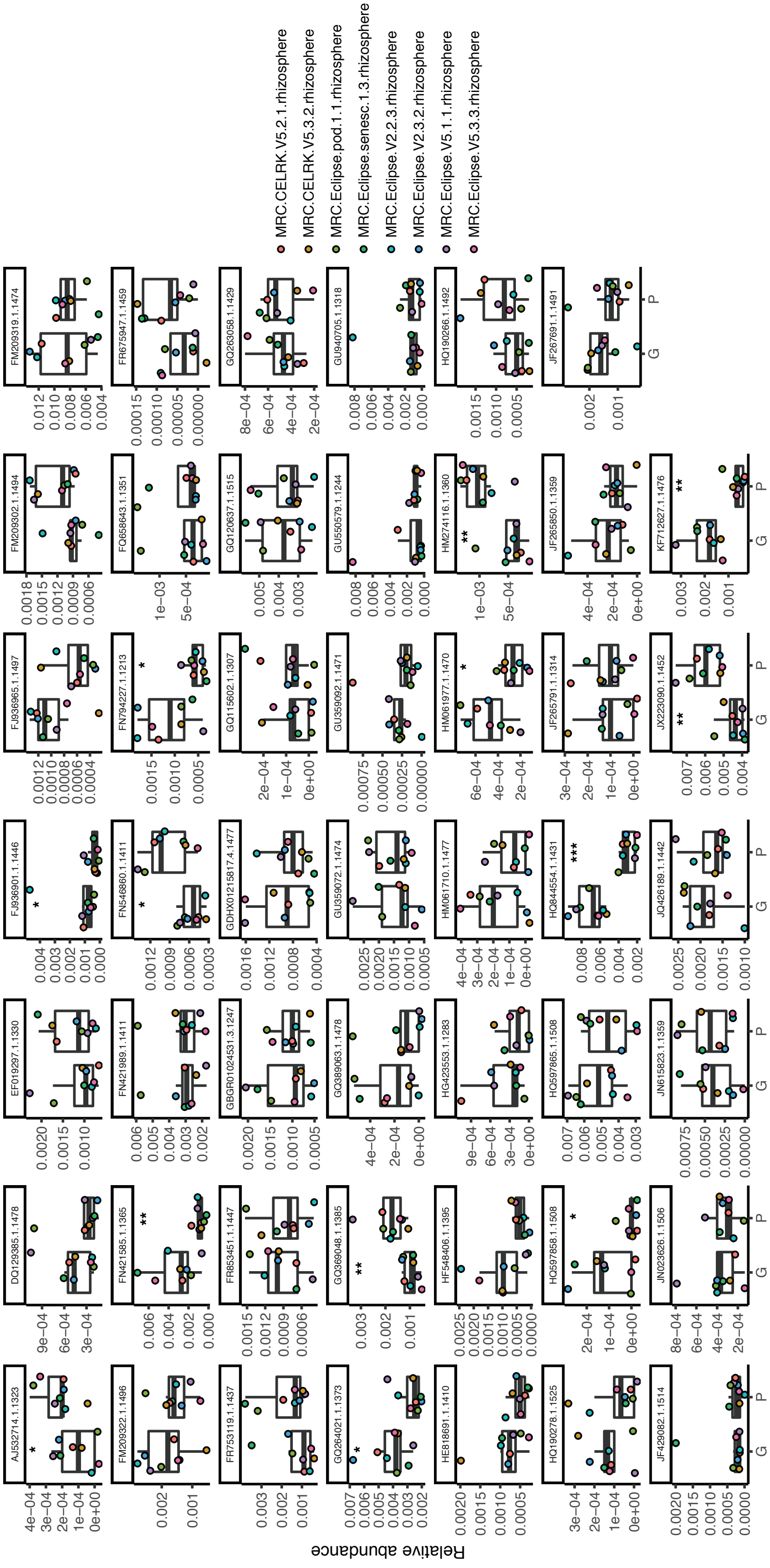


**Fig. S7**: Comparison of relative abundance of the 48 core OTUs between the DNA isolation methods used in the development study (G=Griffith, P=PowerSoil). Statistical difference, determined by Wilcoxon test, is represented as star symbol (*<0.05, **<0.01, ***<0.001). The pints are color coded by sample they derived from.

FigS8

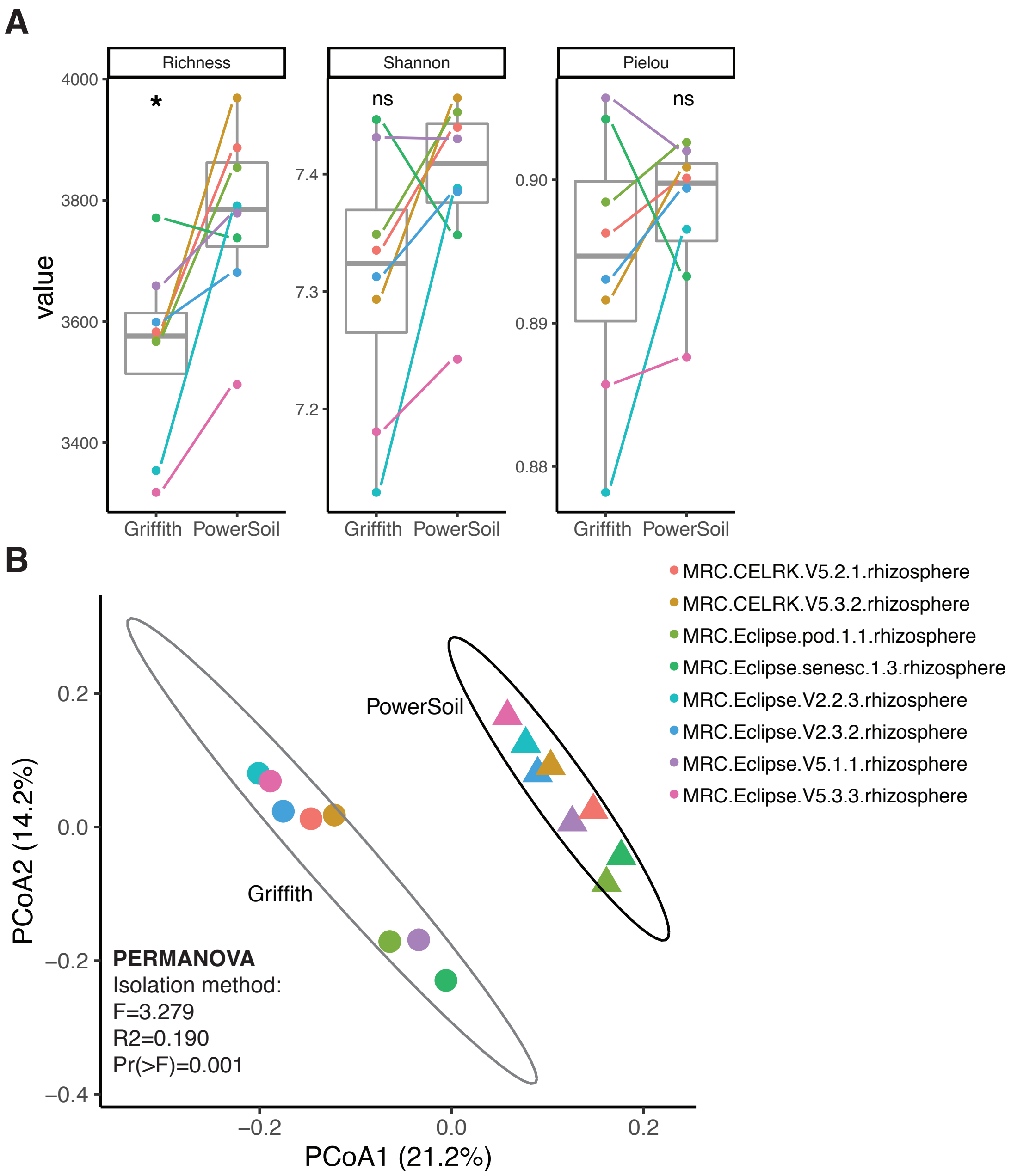


**Fig. S8**: The effect of isolation method on alpha and beta diversity. For alpha diversity richness, Shannon and Pielou indices are presented (A). For the principal coordinates analysis, Bray-Curtis distance matrix was used. Symbols are colored by samples. Wilcoxon test was used to determine statistical differences between isolation methods for the alpha diversity metrices (*<0.05). PERMANOVA was used to determine the effect of isolation method on community structure.
