## Supplementary material for "Prioritizing persistent microbiome members in the common bean rhizosphere: an integrated analysis of space, time, and plant genotype": Table S1

**Table S1:** PERMANOVA results for the 16S rRNA and ITS data. Highly correlated or/and statistical significant values are highlighted in bold.

| **Factor** | **PERMANOVA** | | | |
|---|---|---|---|---|
| | **16S rRNA** | | **ITS** | |
| | $R^2$ | Pr(>F) | $R^2$ | Pr(>F) |
| pH | **0.3708** | **0.001** | **0.2045** | **0.001** |
| sampling location | **0.6643** | **0.001** | **0.6219** | **0.001** |
| genotype | 0.0419 | 0.147 | 0.0337 | 0.213 |
| genotype X sampling location | 0.0300 | 0.114 | 0.0377 | **0.038** |
| irrigation | 0.1205 | **0.001** | 0.1615 | **0.001** |
| fertilization | **0.3550** | **0.001** | **0.3495** | **0.001** |
| soil source (bulk/rhizosphere) | 0.0189 | 0.652 | 0.0161 | 0.809 |
| P | 0.1445 | **0.001** | 0.1244 | **0.001** |
| Nitrogen | 0.1138 | **0.001** | 0.0953 | **0.001** |
| OM | 0.0922 | **0.005** | 0.1414 | **0.001** |
| NO3 | **0.2127** | **0.001** | 0.1151 | **0.002** |
| NH4 | 0.0488 | 0.082 | 0.0586 | **0.026** |
